# Differences in stimulus evoked electroencephalographic entropy reduction distinguishes cognitively normal Parkinson’s disease participants from healthy aged-matched controls

**DOI:** 10.1101/2025.08.06.669023

**Authors:** David T.J. Liley

## Abstract

Parkinson’s disease (PD) is a common neurodegenerative disease best known for its defining motor symptoms. However, it is also associated with significant cognitive impairment at all stages of the disease, with many patients eventually progressing to dementia. Therefore, there exists a significant need to identify objective functional biomarkers that better predict and monitor cognitive decline. While methods that analyse either spontaneous or evoked EEG, due to increasing practical usability and ostensible objectivity, have been investigated, current approaches are limited in that the associated measures are, in the absence of a theoretical basis, purely correlative. To address this shortcoming, we propose calculating changes in evoked EEG amplitude variability, quantified using information theoretic differential entropy (DE), during a three-level passive auditory oddball task, as it is argued this will directly index functional changes in cognition. We therefore estimate changes in stimulus-evoked DE in cognitively normal PD participants (n = 25), both on and off their medication, and in healthy age-matched controls (n = 25), and find substantial stimulus (standard, target, novel) and group differences. Notably, we find the time-course of the return of post-stimulus reductions in DE (i.e., information processing) to pre-stimulus levels delayed in PD compared to healthy controls, thus mirroring the assumed bradyphrenia. The observed changes in DE, together with the corollary increases in resting alpha (8 – 13 Hz) band activity seen in PD, are explained in the context of a well-known macroscopic theory of mammalian electrocortical activity, in terms of reduced tonic thalamo-cortical drive. This method of task-evoked DE EEG amplitude variability is expected to generalise to any situation where the objective determination of cognitive function is sought.

## 1. Introduction

Parkinson’s disease (PD) is best understood as a progressive neurodegenerative disorder affecting movement, its cardinal diagnostic signs being bradykinesia, rigidity, resting tremor and postural instability (Kalia & Lang, 2015). However, it is well known to be associated with cognitive abnormalities at all stages of the disease (Aarsland et al., 2021). As cognitive abnormality may herald disease onset and, at all stages, contribute significant co-morbid suffering it is essential that efforts are made to sensitively identify its presence such that efforts aimed at its management can be optimised.

The most obvious approach to identifying cognitive impairment is through short- and long-form neuropsychological evaluation. However, such methods, nominally regarded as ‘gold standard’, are either too coarse to provide meaningful functional insights or too time consuming to be routinely and repeatedly utilised. For these reasons considerable efforts have been expended in developing alternative metrics of cognitive abnormality that are objective, sensitive, specific and expeditious (Babiloni et al., 2020; Sabbagh et al., 2020; Liu et al., 2023; Risacher & Apostolova, 2023; Liao et al., 2024; Liao et al., 2025). Of those, arguably methods involving recording and analysing the electroencephalogram (EEG) show the most promise and relevance.

Just as blood flow is the *sine qua non* of cardiovascular function, so cortical electrochemical activity is the *sine qua non* of cognitive activity. The absence of cortical electrochemical activity unquestionably implies, at least scientifically, the absence of cognition. At present the best, and possibly forever the only, way of recording such cortical electrochemical activity non-invasively is through EEG and/or magnetoencephalography (MEG). Of these EEG has the greatest likelihood of utility, due to the availability of low-cost hardware and minimal constraints on the recording environment.

Alterations in both resting (spontaneous) and time-locked (evoked, induced) EEG activity have been observed at all stages of the Parkinson’s disease process. Typical resting-state changes include slower alpha band (8 – 13 Hz) activity (Stoffers et al., 2007; Yassine et al., 2024), increases and decreases in eyes-closed alpha band power (Han et al., 2013; Yi et al., 2017; Bin Yoo et al., 2018; Babiloni et al., 2019; Polverino et al., 2022; Gimenez-Aparisi et al., 2023; McKeown et al., 2024; Yassine et al., 2024) and attenuated eyes-open/eyes-closed reactivity (Bosboom et al., 2006; Gimenez-Aparisi et al., 2023; Babiloni et al., 2024).

Changes have also been noted in stimulus evoked (time-locked) EEG activity as well, for example the attenuation, enhancement and delay of P3a and P3b P300 components elicited through auditory or visual oddball tasks (Lagopoulos et al., 1998; Stanzione et al., 1998; Tanaka et al., 2000; Wang et al., 2000; Han et al., 2013; Georgiev et al., 2015; Ozmus et al., 2017; Uslu et al., 2020; Xu et al., 2022).

While such EEG changes point to underlying functional abnormalities, in the absence of any theoretical framework explicitly linking such activity with cognitive function, they are inevitably ‘correlative’ and as such are not direct measures of cognitive function. Can we infer electroencephalographic measures of cognitive function that have a more meaningful theoretical precedence? While this might initially appear to be a hopelessly complicated problem, significant and useful progress can made by asserting an equivalence between information processing and cognition.

Arguably the founding principle of cognitive science is that cognition is brain information processing. Therefore, detecting changes in brain information processing are equivalent to detecting changes in cognition. Because information is operationally defined as the reduction in uncertainty, by quantifying changes in the uncertainty of the EEG response to multiple repeated stimuli, and subsequently looking at the corresponding spatial and temporal interdependencies of such changes, we can expect to obtain a meaningful representation of the informational architecture of the brain as it relates to cognition.

Analogically speaking information flow is to cognition as blood flow is to cardiovascular function.

We apply this approach to EEG recorded in cognitively normal PD participants and a group of age and sex matched cognitively normal controls during the performance of a three-level passive auditory oddball task. Remarkably, we find that at the population level that we can well distinguish the cognitively unimpaired PD participants from their cognitively normal controls. This implies that there are latent functional changes in PD that may have diagnostic and prognostic utility as well as providing a basis for optimising disease management through ongoing monitoring. Finally, we interpret these changes in terms of what is known regarding the phenomenology of PD and attempt to explain them ‘pathodynamically’ in the context of recent theoretical attempts that account for the dynamical genesis of electrocortical activity.

## 2. Methods

### 2.1 Participant EEG data

We choose to reanalyse an EEG data set initially collected to study EEG habituation to novel sensory events (Cavanagh et al., 2018) and made publicly available through the OpenNeuro /NEMAR data repository (https://doi.org/10.18112/openneuro.ds003490.v1.1.0). This data set contains the EEG recorded in PD participants (n = 25), on and off their medication, and in age and sex matched cognitively normal controls (n = 25), while performing a three-level passive auditory oddball task. Parkinson’s and control participants did not differ on any measurements of cognitive function (MMSE), education or pre-morbid intelligence.

The three-level passive auditory oddball task involved participants listening to pseudo random, binaurally presented (external stereo speakers), sequences of standard tones (440 Hz sinusoidal tones, 70% of trials), target tones (660 Hz sinusoidal tones, 15% of trials) and novel distractor sounds (unique sections from a naturalistic sounds database). All tones were presented for 200 ms with a random inter-trial-interval of 500 – 1450 ms. Participants were instructed to count the number of targets and ignore the standards and novel stimuli in two blocks of 100 stimuli, with participants reporting target counts at the end of each block. Participants were free to perform the task with either the eyes open or eyes closed.

During the auditory oddball task 64 channel EEG (extended 10-20 electrode placement, Brain Vision EEG system) was recorded at a sampling frequency of 500 Hz. Prior to the performance of the auditory task participants had 2 min each of resting eyes-closed (EC) and eyes-open (EO) EEG recorded. See Cavanagh et al (2018) for further details regarding participants and the auditory task.

### 2.2 EEG pre-processing and analysis

Continuous EEG was bandpass filtered (1 – 40 Hz, 3^rd^ order Butterworth) and epoched (−500 – 1500 ms) about stimulus onset. Independent Components Analysis was then applied to remove eye-blinks and any obvious ECG artefact, before surface (spherical spline) Laplacian re-referencing. No pre-stimulus baseline correction was applied.

Empirical amplitude (*x*) probability densities, 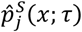, were then estimated (histogram using the Freedman-Diaconis rule for bin width) as a function of epoch stimulus latency, *τ* = [−200 1500] ms, for each electrode, *j*, and stimulus type, *S* ∈ {standard, target, novel}. Inter-trial amplitude standard deviation, 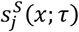, was also calculated.

Temporal changes (i.e., as a function of stimulus latency, *τ*) in the certainty/uncertainty of the time-locked (evoked) response, for a given stimulus and electrode, were estimated in terms of the differential entropy, 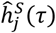, i.e.,

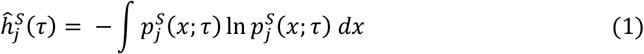

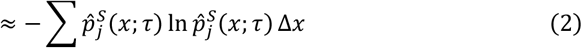

Spectral analysis of resting EO and EC EEG was performed group, channel and subject-wise on Hanning windowed 2 s 50% overlapping segments which were then averaged across group participants, electrodes and windows (i.e., ∼ 95,000 power spectral densities) for each condition.

Resting EC and EO data was also analysed using fixed-order Auto-Regressive Moving-Average (ARMA) modelling to better define EC/EO state differences under the assumption that resting EEG is statistically indistinguishable from a filtered linear random process (Stam et al., 1999; Schwilden & Jeleazcov, 2002; Jeleazcov & Schwilden, 2003; Jeleazcov et al., 2005; Stam, 2005). ARMA modelling enables identification of the dominant noise driven linear relaxation (i.e., stable) oscillatory processes in terms of their frequency and damping (decay rate). Specifically, for a discretely sampled EEG time series *y*[*n*], it is assumed that for stationary intervals it can be well described in terms of the following difference equation

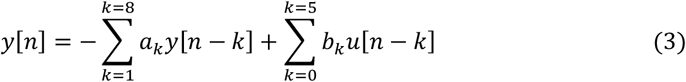

where *a*_*k*_, *b*_*k*_ are the AR and MA coefficients respectively, *b*_0_ = 1 and *u*[*n*] is an iid sequence of random variables having variance 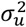. The MA and AR orders (model hyperparameters) of the ARMA model are 8 and 5 respectively, and are derived from the well-known, and physiologically specific, mean field theory of electrocortical activity of Liley et al. (Liley et al., 2002; Liley & Muthukumaraswamy, 2020)

By defining the Z-transform of a discrete time sequence *x*[*n*] as *X* 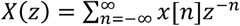 Equation (3) can be rewritten as *A*(*z*)*Y*(*z*) = *B*(*z*)*U*(*z*), where *A*(*z*) = 1 + *a*_1_*z*^−1^ + ⋯ + *a*_8_*z*^−8^ and B(*z*) = 1 + *b*_1_*z*^−1^ + ⋯ + *b*_5_*z*^−5^. Solutions to *A*(*z*) = 0 give the system (filter) poles whereas solutions to *B*(*z*) = 0 give the system (filter) zeros. An estimate of the variance of the innovating random process can be obtained as 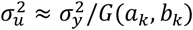, where 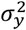 is the variance of the recorded signal and *G*(*a*_*k*_,*b*_*k*_) is the *power gain* of the estimated filter to a unit variance white noise driving process. By convention *σ*_*u*_ is defined as the *cortical input* (CI).

The poles 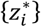, which are solutions to *A*(*z*) = 0, can broadly be thought of as the dominant oscillatory modes that when stochastically forced (i.e., perturbed) and summed approximate the recorded time series *y*[*n*]. Each pole is a complex number and has a corresponding frequency 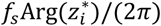 and damping (decay rate) 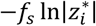, where *f*_s_ is the *sampling frequency*.

To calculate the ARMA coefficients {*a*_*k*_, *b*_*k*_} data was down-sampled to 80 Hz and an (8,5) ARMA model was estimated for 2 s 50% overlapping segments. For each 2 s epoch, CI was calculated and ‘alpha’ poles having frequencies between 5 and 15 Hz were selected in order to calculate their corresponding frequency and damping. For each group, histograms over all participants and electrodes were constructed for CI, alpha pole frequency, and alpha pole damping.

All analyses were performed in MATLAB using the FieldTrip (Oostenveld et al., 2011) analysis toolbox, ARMASA toolbox (Broersen, 2002) and custom scripts.

### 2.3 Statistical analyses

Between group comparisons of the time course of changes in post-stimulus differential entropy were assessed using latency-wise unpaired t-tests together with Bonferroni correction to accommodate for multiple comparisons.

Cluster-based permutation analysis (Nichols & Holmes, 2002; Groppe et al., 2011) was used to identify significant within group differences, as a function of stimulus latency and channel, in post-stimulus differential entropy changes between target and standard conditions. Specifically for each group (Controls, Parkinson’s off medication, Parkinson’s on medication) initial clusters were defined using a paired t-test for alpha = 0.05 (cluster alpha). This was corrected for multiple comparisons by creating a null distribution of the cluster mass (sum absolute t-values) from 10,000 permutations of condition labels and thresholding the original cluster masses at the 95^th^ percentile as the threshold for chance cluster occurrence.

All topographic plots were displayed as the average difference from 450 to 650 ms post-stimulus in differential entropy baseline change between target and standard conditions. False discovery rate corrected significant differences are indicated by red highlighted electrodes. Multiple comparison correction was performed by creating a null distribution of the maximum t-value across all electrodes from 10,000 permutations of the condition labels and thresholding the original t-values at the 95^th^ percentile as the threshold for chance occurrence.

## 3. Results

### 3.1 Differential entropy (DE) varies with stimulus type and latency

We first investigated how evoked variability, as quantified by differential entropy (DE, see Equation 2), changed following stimulus onset. Figure 1 shows the grand average (i.e., across all participants and electrodes) baseline changes in DE for all three participant groups across the three oddball (standards, targets and novel) stimuli. In all participant groups changes induced by standard stimuli from a pre-stimulus baseline were small compared to those induced by target and novel stimuli. In contrast target and novel stimuli induced global reductions in DE that were maximal ∼ 500 ms after stimulus onset, before increasing towards pre-stimulus baseline, and in some cases overshooting before decreasing. In general, post-stimulus reductions in DE induced by targets were much greater than for novel stimuli.

**Figure 1.**
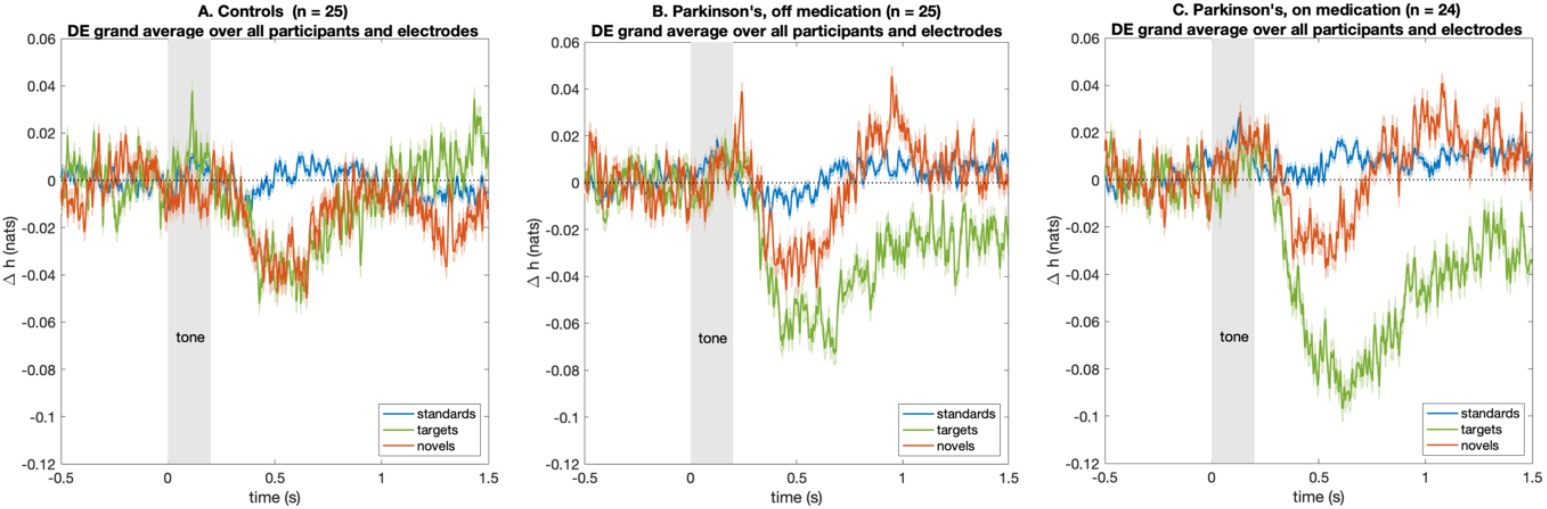
Grand average differential entropy (DE) as a function of stimulus onset latency. A. Change in baseline DE grand average (over all participants and electrodes) for Control (N = 25) participants in response to each type (standard, target, novel) of auditory stimulus. B. Change in baseline DE grand average (over all participants and electrodes) for Parkinson’s participants OFF their medication (N = 25) in response to each type (standard, target, novel) of auditory stimulus. C. Change in baseline DE grand average (over all participants and electrodes) for Parkinson’s participants ON their medication (N = 24) in response to each type (standard, target, novel) of auditory stimulus. Change in DE is quantified in nats (natural unit of information entropy defined in terms of the natural logarithm, see Eq.2). Solid lines represent mean over all respective participants and electrodes with shaded areas the standard error in the mean.

Statistically very similar results (not shown) were obtained if DE was estimated on the basis that inter-trial amplitude was Gaussian i.e.,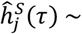 ln 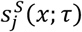. Thus, temporal changes in variability appear to be driven by changes in multiple independent processes rather than a single coupled stochastic system.

### 3.2 Target induced changes in differential entropy (DE) differ across disease and medication status

While changes in DE for standard and novel stimuli were similar across groups, this was not the case for targets. Figure 2 shows that all groups differed significantly in DE as a function of target stimulus latency. Unexpectedly, PD participants both off and on their medication, had greater reductions compared to controls in DE following target onset, as well as a slower return to baseline. Such a reduction in DE was greatest for PD participants *on* their medication, with both PD groups evincing a similar time course of return to baseline.

**Figure 2.**
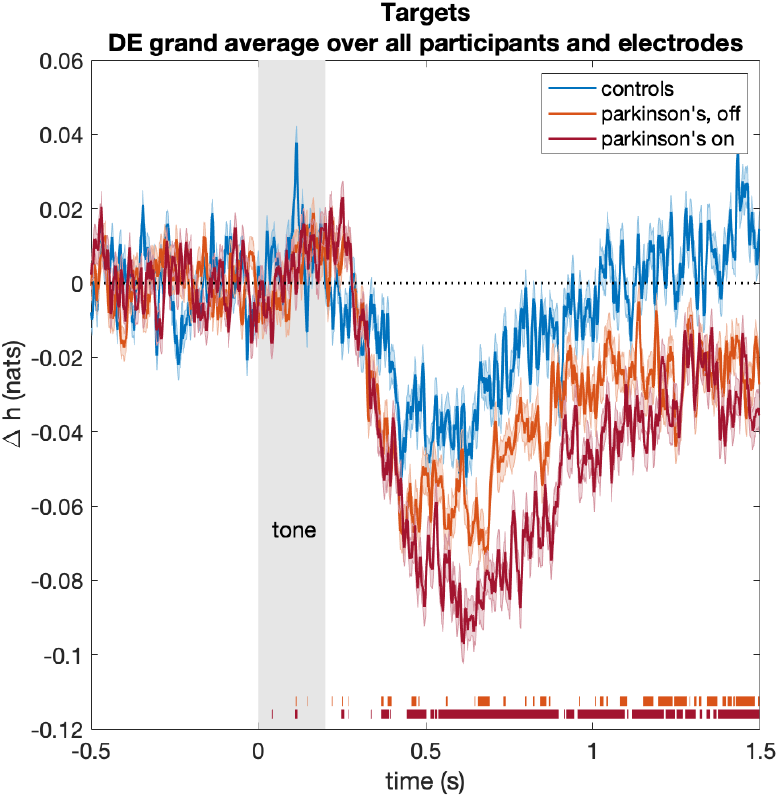
Time course of target induced changes in baseline DE for control and PD participants. Coloured bands at the bottom of the figure indicate false discovery rate corrected (Bonferroni, see section 2.3) significant (*p* < 0.05) group differences (all electrodes and group participants) between: PD participants (off medication, light red) and Control participants, and PD participants (on medication, dark red). Solid lines represent mean over all respective participants and electrodes with shaded areas the standard error in the mean.

As these DE changes have been calculated on a broadband signal we sought to determine what frequency band power decreases were principally driving these DE reductions. By bandpass filtering in the canonical EEG bands of delta (< 4 Hz), theta (4 – 8 Hz), alpha (8 – 13 Hz), and beta (8 – 30 Hz), we recalculated ‘band-limited’ DE and found that it was reductions in alpha band activity that were most highly correlated with changes in ‘broad band’ DE (results not shown).

### 3.3 Topographic changes in differential entropy (DE) are occipito-parietally focused

Given the substantial differences in globally calculated changes in DE we sought to investigate their topographic distribution. In order to minimise the consequences of inter-individual variability we chose to calculate topographic changes in DE between the target and the standard condition. Figure 3 shows the results of permutation analysis applied to the corresponding high density time locked EEG recordings. As can be seen while significant changes between standard and target stimuli are seen in all conditions, such changes are particularly striking for PD on and off their medication. The weak centro-parietal electrode significance in controls, becomes quite spatio-temporally extensive in PD participants off their medication, and even more extensive when PD participants are *on* their medication.

**Figure 3.**
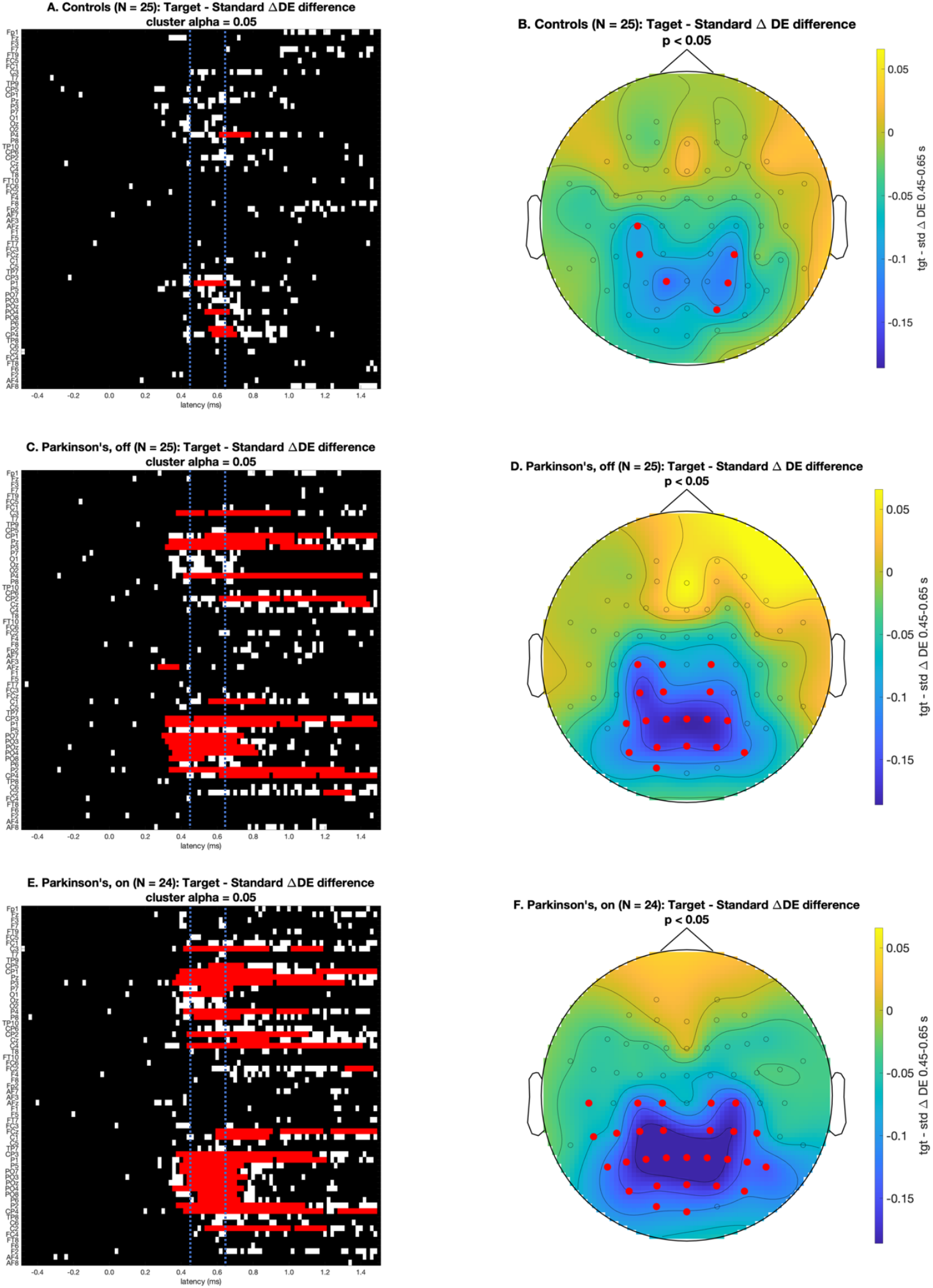
Spatio-temporal differences between target and standard conditions in controls and PD participants. A, C and E: Raster plots (coarse grained to 10 ms) indicating significantly different (red, *p* < 0.05, family wise error rate (FWER) corrected via mass univariate cluster-based permutation test, see section 2.3) univariate temporal clusters between target and standard conditions for controls (A), PD participants off their medication (C) and PD participants on their medication (E). Red squares indicate univariate stimulus latency clusters which are significantly different between target and standard conditions, white squares indicate significant differences (cluster alpha *p* < 0.05) between target and standard conditions before FWER correction and black squares indicating no significant difference. B, D and F: Topographic plots of the target – standard difference in DE averaged over 0.45 - 0.65 s (blue dashed lines; A, C and E) post-stimulus. Highlighted electrodes (red) indicate a significant groupwise difference (*p* < 0.05, FWER corrected via maximum t-statistic permutation test, see section 2.3) between target and standard conditions.

The pattern of these spatio-temporal changes can be used to define a very preliminary functional biomarker. For example, we can choose a set of electrodes (i.e., Pz & P2) that 1) have the largest significant DE difference between target and standard conditions and 2) are not significantly different between target and standard conditions in controls, over a time (latency) of interest of [0.5 0.55] s that represents a midpoint of the region encompassing the largest number of significant electrodes (see Figure 3A, C & E). Doing so reveals an AUC (area under the curve) of the ROC (receiver operating characteristic) curve of 0.745 (figure not shown).

### 3.3 Resting eyes-closed (EC) and eyes-open (EO) alpha EEG is increased in Parkinson’s disease (PD)

Given reductions in alpha (8 – 13 Hz) band power is the principal driver of decreases in post-stimulus DE, and that these post-stimulus changes are noticeably larger in PD than in controls, we wondered whether there were corollary changes in the corresponding resting EEG. Figure 4 shows global periodogram estimates of resting eyes-closed (EC) and eyes-open (EO) EEG activity in control participants and PD participants on and off their medication. Clear differences between control and PD participants are observed: alpha band power is increased in PD.

**Figure 4.**
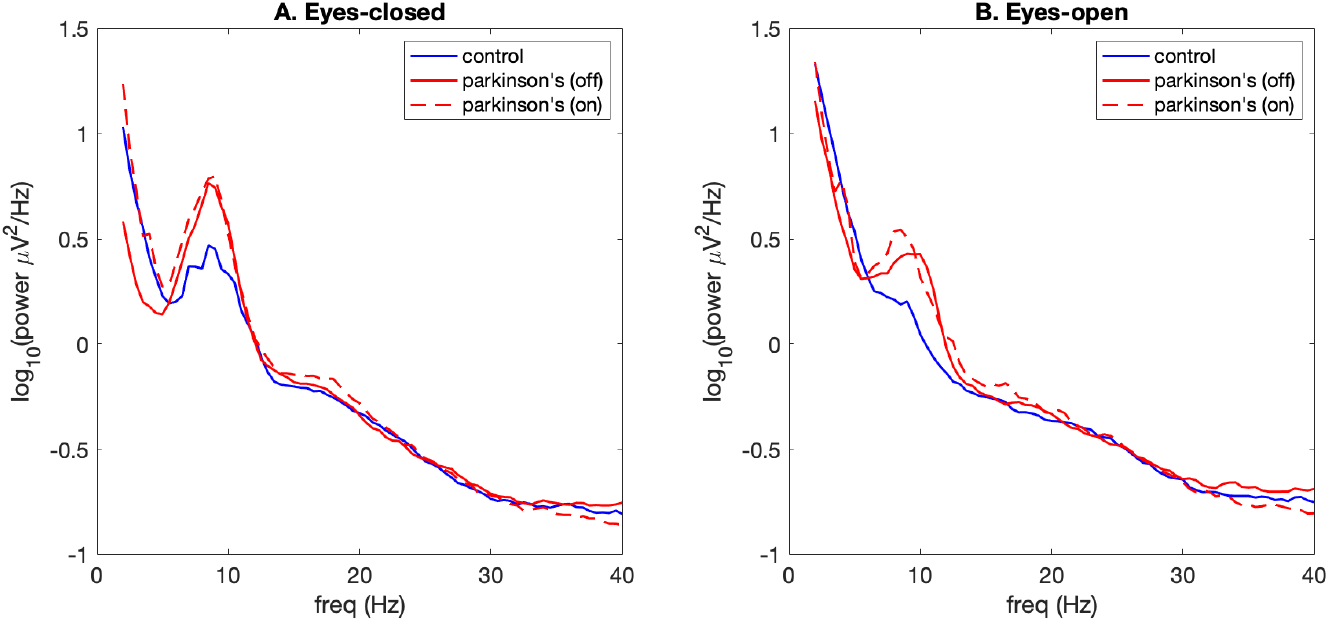
Eyes-closed (EC) and eyes-open (EO) power spectral differences between control and PD participants. A. EC grand average (participants and electrodes) power spectral density for Controls (N = 25), PD off medication (N = 25) and PD on medication (N = 24). Power spectral density calculated on Hanning-windowed 2 s 50% overlapping segments and then averaged over all segments, electrodes and participants. These results should be compared with those of Figure 2a in McKeown et al., 2023, where it is suspected they have inadvertently averaged individual log semi-log transformed spectra such that the average power represents a geometric mean over participants.

To further explore the basis for these power changes we also performed fixed-order auto-regressive moving-average (ARMA) time series modelling (see Methods 2.2) on the resting EEG. ARMA analysis identifies the dominant damped oscillatory modes, in terms of their frequency and decay rate, that when stochastically forced (i.e., randomly perturbed) account for EEG spectral activity, as well as enabling an estimate of the amplitude of the stochastic forcing (*cortical input*). Figure 5 summarises the results of this analysis. Notably, and recapitulating previous results (Liley & Muthukumaraswamy, 2020), the lower amplitude EO activity has higher alpha band damping than the higher amplitude, less weakly damped, EC activity. Accompanying the increase in damping between EC and EO states are increases in alpha pole frequency. The significance of these changes is elaborated in the Discussion.

**Figure 5.**
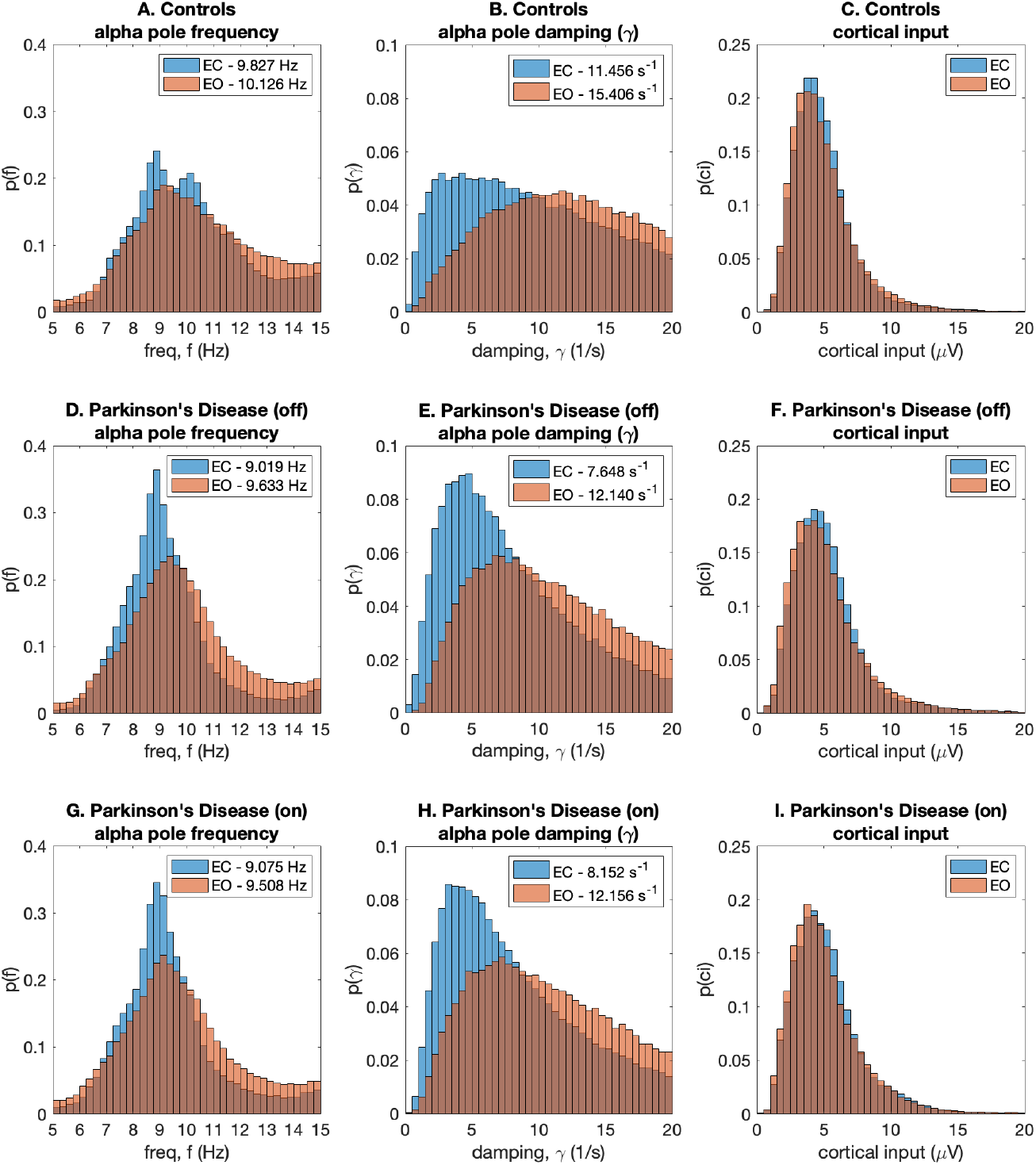
Fixed-order ARMA analysis of EC and EO state: Empirical probability density functions (ePDFs) for ARMA estimated alpha pole frequency (A, D, G), alpha pole damping (B, E, H) and cortical input/driving (C, F, I) across all respective group participants and recording electrodes. Figures in figure legends represent medians. Full range of damping values not shown for ePDFs. See Methods 2.2 for further details regarding fixed-order ARMA modelling.

Remarkably EC and EO alpha pole frequency (Figure 5A, D & G) and damping (Figure 5B, C & E) are reduced in PD participants, both on and off their medication, when compared to controls. In contrast cortical input appears largely unchanged. As argued in the Discussion these changes in resting alpha band activity, and the corresponding changes in DE, can be theoretically accounted for in terms of reductions in thalamo-cortical modulatory input to cortical inhibitory interneurons.

Based on the ARMA modelling we can also calculate a more specific measure of EC → EO reactivity based on differences in damping, rather than changes in band power as is more typically done (see legends in Figure 5B, E & H). Thus EC → EO reactivity for PD (off) 4.492 s^−1^> PD (on), 4.004 s^−1^ ≈ Control, 3.950 s^−1^, contrary to what has been previously reported using measures of power spectral density (Bosboom et al., 2006; Gimenez-Aparisi et al., 2023; Babiloni et al., 2024), where reactivity in PD is impaired.

## 4. Discussion

By characterising stimulus evoked EEG activity in terms of inter-trial variability, specifically differential entropy, we are able to differentiate, at the population level, PD participants (on and off medication) from age-matched controls. Specifically, we found that in controls and PD participants that following the onset of a target stimulus inter-trial variability, as assessed using differential entropy (DE), decreased substantially from the pre-stimulus baseline reaching a maximum reduction at approximately 400 – 700 ms after stimulus onset. It is reasonable to assume that such changes in variability mirror macroscopically the post-stimulus variability quenching observed in the spike trains/counts of single neural activity, as quantified using the Fano Factor on time-averaged neuronal spike trains (Stein et al., 2005; Churchland et al., 2010; Ribeiro et al., 2024). In contrast the reported changes in the time-ensemble event related potentials (ERPs) between groups were quantitatively much less evident (Cavanagh et al., 2018).

Such reductions were larger, more spatially extensive and more sustained in PD participants *off* their medication, and larger and more extensive again when PD participants were *on* their medication. Such post-stimulus reductions in DE were also evident for the novel stimuli, however there appeared little variation over groups. In contrast standard stimuli evoked little change in post-stimulus DE.

In contrast to similar efforts at characterising intertrial variability we did not focus on any assumed ERP components (e.g., P3a, P3b, etc.)(Ouyang et al., 2017; Depuydt et al., 2023) and nor did we characterise directly changes in relative or absolute intertrial variance as a function of stimulus latency (Arazi, Censor, et al., 2017; Arazi, Gonen-Yaacovi, et al., 2017; Arazi et al., 2019; Daniel et al., 2019; Wolff et al., 2019). Instead, we chose to calculate the intertrial variability in terms of DE as a function of stimulus latency, such that our measure of variability was distribution free and thus as general as possible, and to make specific an information theoretic link. We can thus interpret the observed electroencephalographic reductions in uncertainty informationally, with greater post-stimulus reductions in DE (or some time integrated DE) implying a more informative stimulus. Therefore, as required by the task target stimuli were more informative than novel stimuli, which were more informative than standard stimuli. Further, the delayed return of target DE to baseline in PD participants would seem to imply slower information processing and thus be expected to correlate with the assumed bradyphrenia.

### 4.1 Significance of increased DE reductions in PD

Unexpectedly target stimuli were more informative in PD, both on and off their medication, when compared to the age-matched controls. This is a surprising result as it suggests that perceptual sensitivity to the auditory stimuli, and possibly other types of sensory stimuli, may be enhanced in PD, contrary to the perceptual impairments that are typically reported (Bodis-Wollner, 2003; Cecchini et al., 2015; Weil et al., 2016; Nieto-Escamez et al., 2023). It is therefore interesting that enhancements in auditory perception in PD have been reported. For example, PD individuals when asked to produce speech with normal loudness (as judged by an attending speech pathologist) perceive themselves as producing abnormally loud speech or shouting (Fox & Raming, 1997). Similarly, when asked to estimate loudness intensity PD participants have been reported to estimate greater intensities when compared to healthy control participants (Ho et al., 1999). Further, it has been reported that PD individuals have an enhanced ability to detect vocal pitch differences compared to control participants, during a production condition, where participants were required to identify shifts in their vocal pitch that was shifted in real time (Mollaei et al., 2019).

### 4.2 Differences between PD and controls can be explained by a mean field theory of resting alpha EEG

The increased resting alpha activity (Figure 4), and the reduced frequency (Figures 5 A, D & G), observed in PD individuals compared to control participants recapitulates results previously reported involving the reanalysis of multiple resting state datasets that include many hundreds of PD and control participants (Norouzi et al., 2025). The fact that such changes are the result of reductions in the damping (Figure 5 B, E & H) and frequency of stochastically-forced linear alpha band oscillatory activity insinuates a role for mean field theories of macroscopic electrocortical activity in accounting for the identified group differences (Coombes, 2005; Deco et al., 2008; Cook et al., 2022; Glomb et al., 2022; Bastiaens et al., 2025). Mean field theories, also referred to as neuronal population or neural mass action models, aim to mathematically account for macroscopic cortical brain dynamics (most particularly the EEG) by assuming it is sufficient to account for the mean activity of interacting excitatory and inhibitory neuronal populations, each comprising many thousands of neurons. The majority of the models developed to date (see Bastiaens et al., 2025 for a recent review) are rate-based (i.e., spiking behaviour is not explicitly modelled) and focus on accounting for the dynamical genesis and features of the mammalian alpha rhythm, based on reasonable and parsimonious assumptions regarding cortical anatomy and physiology.

Of these mean field models, that of Liley et al. (1999,2002) is of particular note in that it is physiologically quite detailed such that biological meaningful inferences can be made from parametric changes. This model accounts for the resting alpha rhythm in terms of a broadband noise synaptic input (arising cortically and subcortically) driving reverberant local activity arising from feedforward and feedback inhibitory and excitatory neuronal population connectivity. Mathematically speaking, white noise drives a stable linear alpha band damped resonance. Detailed efforts to fit this model to resting EC and EO EEG data have revealed that the parameters associated with neural inhibition are the most sensitive determinants of model alpha band dynamics (Hartoyo et al., 2019, 2020). On this basis it has been hypothesised that changes in a single model parameter, *p*_*ei*_, corresponding to the strength of tonic, thalamo-cortical excitatory input to cortical inhibitory neurons, is sufficient to explain changes in alpha band power between EC and EO states, and by inference other states (Hartoyo et al., 2020). The effect of increases/decreases in this model parameter is to increase/decrease the damping (i.e., decay rate), and decrease/increase the frequency, of stochastically forced alpha band oscillatory activity.

There is evidence to suggest that thalamo-cortical (excitatory) drive, particularly to motor cortex, is reduced in PD as a direct consequence of elevated basal ganglia inhibition (Galvan et al., 2015; McGregor & Nelson, 2019; Chen et al., 2023). On this basis we assert that reductions in *p*_*ei*_ in PD not only account for the observed increased alpha band activity, and reduced frequency, but also argue that it leads to the identified changes in stimulus evoked DE reduction. Figure 6 represents a schematic attempt at such an argument. The essence of the argument is that reducing *p*_*ei*_ increases the sensitivity of alpha band damping (which determines resting alpha power) to changes in cortico-cortical excitatory input to cortical inhibitory neurons that arise as a consequence of stimulus induced cortical activity. On this basis stimulus mediated increases in cortical excitatory activity induce greater reductions in alpha band activity (the major driver of decreases in DE) when *p*_*ei*_ is diminished in PD.

**Figure 6.**
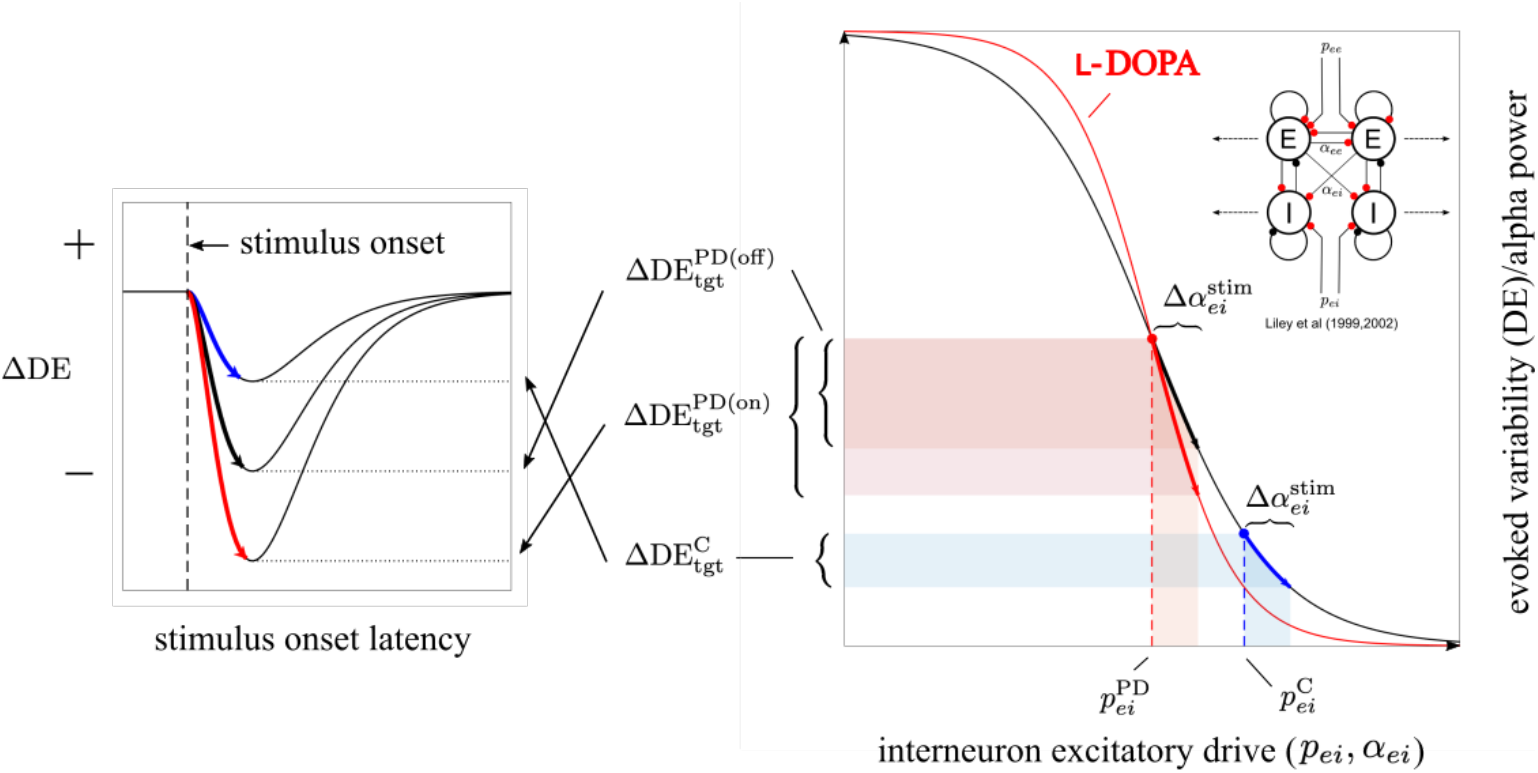
Differences between participant groups in differential entropy changes to targets might be theoretically explicable in terms of a neural population model of the alpha rhythm. Neural field/population models offer a promising approach to understanding the dynamical genesis of the resting alpha rhythm. In particular, the model of Liley et al (Liley et al., 1999; Liley et al., 2002) claims to account for the alpha rhythm and its dynamical genesis in terms of a physiologically and anatomically well parametrised mass action framework. It theoretically posits that the alpha rhythm arises from reverberant activity between interconnected populations of cortical excitatory and inhibitory (inter-) neurons (top right inset, excitatory synaptic activity = red, inhibitory synaptic activity = black), and that changes in its amplitude are driven by changes in system level damping, and not changes in ‘neural synchrony’ as is widely assumed. Detailed investigations involving fitting the model to empirical eyes-open (EO) and eyes-closed (EC) EEG data has revealed that just a single parameter, *p*_*ei*_, corresponding to the strength of a tonic, extra-cortical (thalamic) excitatory input to the cortical inhibitory neuronal population, is sufficient to account for the reduction in alpha power between EC and EO conditions (Hartoyo et al., 2020). Increases in *p*_*ei*_ are theoretically associated with an increase in system level damping and a corresponding reduction in the amplitude of alpha oscillations. If we hypothesise that the magnitude of this tonic thalamo-cortical drive is reduced in Parkinson’s disease (for which there seems to exist empirical evidence in animal models e.g. Chen et al., 2023; Galvan et al., 2015) then we can broadly account for the observed group differences in the time course of post-stimulus changes in intertrial EEG variability (i.e., differential entropy - DE). To see this, consider the sigmoid curve (black line) in the right-hand figure which schematically details the theoretical relationship between alpha power and *p*_*ei*_. A sigmoid curve can be seen as reasonable as we expect alpha power to be bounded above by a maximum value and below by zero. Increases in *p*_*ei*_ attenuate the amplitude of resting alpha, whereas decreases increase the amplitude of alpha. Thus if *p*_*ei*_ is decreased in Parkinson’s disease then we expect alpha to be enhanced – which is what we observe (see Figure 4.). However, a reduction in *p*_*ei*_ (i.e., from 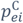 to 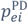) shifts us to a steeper part of the sigmoid curve, such that any subsequent perturbation in interneuron excitatory drive, such as due to phasic excitatory input from other areas of cortex (via long range cortico-cortical axons) occurring in response to target stimulus activity (i.e.,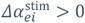), will be associated with greater changes (reductions) in alpha power/variability (i.e.,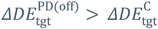), than for equivalent perturbations about larger (normal) values of *p*_*ei*_ (i.e.,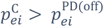). It has been assumed that reductions in the intertrial variability are being principally driven by the attenuation of alpha band activity (which is what we find when we bandpass filter the data into the canonical EEG frequency bands and assess the magnitude of the correlations between the broadband stimulus induced changes in DE and the corresponding ‘narrow band’ DE changes; results not shown). We posit that dopaminergic medication will act to modulate the sigmoidal relationship between *p*_*ei*_ and alpha band power but otherwise not alter tonic *p*_*ei*_. Specifically, and in order to phenomenologically account for our results, we assert that dopaminergic medication increases the slope of this sigmoid (red curve in right-hand figure) such that changes in stimulus related phasic excitatory activity (i.e.,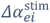) induce greater reductions in alpha band power/variability 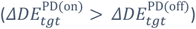, as indicated by the red line segment, for equivalent perturbations in 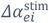. We have ignored the effect of any changes in *p*_*ee*_ and 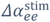 that will necessarily accompany changes in *p*_*ei*_ and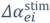, even though they share the same cortical (pyramidal neurons) and subcortical (thalamic) sources, for the reason that theoretical analysis implies that effect of reducing *p*_*ei*_ is considerably stronger in increasing alpha band power than reductions in *p*_*ee*_ are at reducing it (Hartoyo et al., 2020). Abbreviations/Symbols: C, control/normal; DE, differential entropy (see Equation 2); PD, Parkinson’s disease; *α*_*ee*_, cortico-cortical (excitatory) input to excitatory (pyramidal) neurons; *α*_*ei*_, cortico-cortical (excitatory) input to inhibitory (inter-) neurons;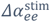, maximum stimulus induced change in 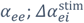, maximum stimulus induced change in *α*_*ei*_; *p*_*ee*_, thalamo-cortical input to excitatory (pyramidal) neurons; *p*_*ei*_, thalamo-cortical input to inhibitory (inter-) neurons;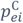, ‘resting’ *p*_*ei*_ in controls/normal condition;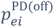, ‘resting’ *p*_*ei*_ in PD; *Δ*DE, change in DE from pre-stimulus baseline;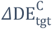, ‘peak’ *ΔDE* due to target stimulus in controls/normals;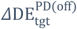, ‘peak’ *ΔDE* due to target stimulus in PD off medication; 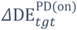’peak’ *Δ*DE due to target stimulus in PD on medication.

### 4.3 Relationship of DE to time-frequency methods

We chose to quantify across trial variability using DE, but a little thought will reveal that such a calculation is broadly equivalent to calculating across-trial amplitude variability in terms of either standard deviation, variance, or within-trial broadband ‘instantaneous’ root-mean-square (RMS) amplitude or broadband ‘instantaneous’ total spectral power, with the latter two not meaningful calculable given the trade-offs of time and frequency resolution. Nevertheless, by accepting limitations on resolution, narrow band absolute or relative estimates of the latter are often calculated with the attendant changes been referred to as generally as either Event-Related Spectral Perturbation (ERSP), or more specifically as Event-Related Desynchronization (ERD) for power decreases, or Event-Related Synchronisation (ERS) for power increases. While such time-frequency decompositions have become part of the standard methodological armamentarium in cognitive electrophysiology, it can be argued on the basis of our results that they obscure direct informational interpretations in favour of second-order oscillatory exegeses.

### 4.4 ERP vs DE

How does our time-locked method accord with the time-ensemble average of the classically defined cortical ERP? It is not often appreciated but the middle and late cognitive ERP is essentially a theoretical construct, based on a central premise that there is an invariant response embedded in a sea of uncorrelated electroencephalographic noise. This has led to the oft mentioned limitation of EEG in that it is a ‘low signal-to-noise’ measurement. In contrast the assessment of inter-trial amplitude variability, in terms of DE, does not depend on an underlying model.

Further, while not often explicitly stated this model implies that over the assumed time course of the ERP that the additive EEG will be *homoscedastic* i.e., the variance is constant.

But as we (Figures 1 & 2) and others (Arazi, Censor, et al., 2017; Arazi, Gonen-Yaacovi, et al., 2017; Arazi et al., 2019; Daniel et al., 2019; Wolff et al., 2019) have demonstrated this is clearly not the case.

### 4.5 Implications for the Free Energy Principle

As much as it can be understood the *Free Energy Principle* (FEP)(Friston, 2010) seems to suggest that cognition and behaviour centrally depend on the minimisation of *surprisal* i.e., minimizing the ‘surprise’ of a sensory or behavioural event. In an attempt to neurophysiologically operationalise such a hypothesis an ‘phenomenal’ appeal (Garrido et al., 2009; Friston, 2010) is made to the time ensemble averaged oddball event related potential (ERP), in which the infrequent random stimulus induces an MMN/P3a/P3b response which is assumed to correspond to a prediction error. However, the basis for this hypothesis, in the context of our results would seem problematic.

A less variable response (i.e., lower DE) to a stimulus would imply it is more predictable, whereas a more variable response (i.e., higher DE) to a stimulus would imply it is less predictable. Thus, according the FEP the more frequent standard stimulus should be associated with a lower prediction error than the less frequent target stimulus. However, we observe the *complete opposite*: the time integrated DE for the target is *less* than the time integrated DE for the standard.

## 5. Conclusion

It has been demonstrated that PD participants can be differentiated from healthy aged-matched controls at the population level in terms of changes in evoked EEG variability as quantified using information theoretic DE. Further, it has been argued that such changes can be explained theoretically in the context of a physiologically detailed mean field theory of electrocortical activity when the corollary changes in resting alpha band activity are taken into account. The changes in evoked DE in PD participants may eventually serve as a useful objective longitudinal measure of cognitive function and integrity that may help better identify and manage the accompanying cognitive impairment. Indeed, a simply constructed index involving only a pair of electrodes and a short post-stimulus time of interest already exhibits a useful ROC curve AUC of 0.75, a value that is expected to be enhanced when more detailed analyses are performed that drill-down to quantify regional variations and inter-regional correlations.

The utility of quantifying changes in evoked variability may extend to measuring cognitive function in any situation where objective information regarding cognition is sought. Indeed, DE and other related measures may find use as ‘features’ for training AI/ML architectures for disease diagnosis and monitoring.

## References

Aarsland, D., Batzu, L., Halliday, G. M., Geurtsen, G. J., Ballard, C., Ray Chaudhuri, K., & Weintraub, D. (2021). Parkinson disease-associated cognitive impairment. Nat Rev Dis Primers, 7(1), 47. 10.1038/s41572-021-00280-3

Arazi, A., Censor, N., & Dinstein, I. (2017). Neural Variability Quenching Predicts Individual Perceptual Abilities. J Neurosci, 37(1), 97–109. 10.1523/JNEUROSCI.1671-16.2016

Arazi, A., Gonen-Yaacovi, G., & Dinstein, I. (2017). The Magnitude of Trial-By-Trial Neural Variability Is Reproducible over Time and across Tasks in Humans. eNeuro, 4(6). 10.1523/ENEURO.0292-17.2017

Arazi, A., Yeshurun, Y., & Dinstein, I. (2019). Neural Variability Is Quenched by Attention. J Neurosci, 39(30), 5975–5985. 10.1523/JNEUROSCI.0355-19.2019

Babiloni, C., Blinowska, K., Bonanni, L., Cichocki, A., De Haan, W., Del Percio, C., Dubois, B., Escudero, J., Fernandez, A., Frisoni, G., Guntekin, B., Hajos, M., Hampel, H., Ifeachor, E., Kilborn, K., Kumar, S., Johnsen, K., Johannsson, M., Jeong, J., … Randall, F. (2020). What electrophysiology tells us about Alzheimer’s disease: a window into the synchronization and connectivity of brain neurons. Neurobiol Aging, 85, 58–73. 10.1016/j.neurobiolaging.2019.09.008

Babiloni, C., Del Percio, C., Lizio, R., Noce, G., Lopez, S., Soricelli, A., Ferri, R., Pascarelli, M. T., Catania, V., Nobili, F., Arnaldi, D., Fama, F., Orzi, F., Buttinelli, C., Giubilei, F., Bonanni, L., Franciotti, R., Onofrj, M., Stirpe, P., … De Pandis, M. F. (2019). Levodopa may affect cortical excitability in Parkinson’s disease patients with cognitive deficits as revealed by reduced activity of cortical sources of resting state electroencephalographic rhythms. Neurobiol Aging, 73, 9–20. 10.1016/j.neurobiolaging.2018.08.010

Babiloni, C., Noce, G., Tucci, F., Jakhar, D., Ferri, R., Panerai, S., Catania, V., Soricelli, A., Salvatore, M., Nobili, F., Arnaldi, D., Fama, F., Buttinelli, C., Giubilei, F., Onofrj, M., Stocchi, F., Vacca, L., Radicati, F., Fuhr, P., … Del Percio, C. (2024). Poor reactivity of posterior electroencephalographic alpha rhythms during the eyes open condition in patients with dementia due to Parkinson’s disease. Neurobiol Aging, 135, 1–14. 10.1016/j.neurobiolaging.2023.11.010

Bastiaens, S. P., Momi, D., & Griffiths, J. D. (2025). A comprehensive investigation of intracortical and corticothalamic models of the alpha rhythm. PLoS Comput Biol, 21(4), e1012926. 10.1371/journal.pcbi.1012926

Bin Yoo, H., Concha, E. O., De Ridder, D., Pickut, B. A., & Vanneste, S. (2018). The Functional Alterations in Top-Down Attention Streams of Parkinson’s disease Measured by EEG. Sci Rep, 8(1), 10609. 10.1038/s41598-018-29036-y

Bodis-Wollner, I. (2003). Neuropsychological and perceptual defects in Parkinson’s disease. Parkinsonism Relat Disord, 9 Suppl 2, S83–89. 10.1016/s1353-8020(03)00022-1

Bosboom, J. L., Stoffers, D., Stam, C. J., van Dijk, B. W., Verbunt, J., Berendse, H. W., & Wolters, E. (2006). Resting state oscillatory brain dynamics in Parkinson’s disease: an MEG study. Clin Neurophysiol, 117(11), 2521–2531. 10.1016/j.clinph.2006.06.720

Broersen, P. (2002). Automatic spectral analysis with time series models. IEEE Trans Instrum Meas, 51, 211–216. 10.1109/19.997814

Cavanagh, J. F., Kumar, P., Mueller, A. A., Richardson, S. P., & Mueen, A. (2018). Diminished EEG habituation to novel events effectively classifies Parkinson’s patients. Clin Neurophysiol, 129(2), 409–418. 10.1016/j.clinph.2017.11.023

Cecchini, M. P., Fasano, A., Boschi, F., Osculati, F., & Tinazzi, M. (2015). Taste in Parkinson’s disease. J Neurol, 262(4), 806–813. 10.1007/s00415-014-7518-1

Chen, L., Daniels, S., Dvorak, R., & Chu, H. Y. (2023). Reduced thalamic excitation to motor cortical pyramidal tract neurons in parkinsonism. Sci Adv, 9(34), eadg3038. 10.1126/sciadv.adg3038

Churchland, M. M., Yu, B. M., Cunningham, J. P., Sugrue, L. P., Cohen, M. R., Corrado, G. S.,Newsome, W. T., Clark, A. M., Hosseini, P., Scott, B. B., Bradley, D. C., Smith, M. A., Kohn, A.,Movshon, J. A., Armstrong, K. M., Moore, T., Chang, S. W., Snyder, L. H., Lisberger, S. G., … Shenoy, K. V. (2010). Stimulus onset quenches neural variability: a widespread cortical phenomenon. Nat Neurosci, 13(3), 369–378. 10.1038/nn.2501

Cook, B. J., Peterseon, A. D. H., Woldman, W., & Terry, J. R. (2022). Neural Field Models: a mathematical overview and unifying framework. Math Neuro Appl, 2. 10.46298/mna.7284

Coombes, S. (2005). Waves, bumps, and patterns in neural field theories. Biol Cybern, 93(2), 91–108. 10.1007/s00422-005-0574-y

Daniel, E., Meindertsma, T., Arazi, A., Donner, T. H., & Dinstein, I. (2019). The Relationship between Trial-by-Trial Variability and Oscillations of Cortical Population Activity. Sci Rep, 9(1), 16901. 10.1038/s41598-019-53270-7

Deco, G., Jirsa, V. K., Robinson, P. A., Breakspear, M., & Friston, K. (2008). The dynamic brain: from spiking neurons to neural masses and cortical fields. PLoS Comput Biol, 4(8), e1000092. 10.1371/journal.pcbi.1000092

Depuydt, E., Criel, Y., De Letter, M., & van Mierlo, P. (2023). Single-trial ERP Quantification Using Neural Networks. Brain Topogr, 36(6), 767–790. 10.1007/s10548-023-00991-8

Fox, C. M., & Raming, L. O. (1997). Vocal sound pressure level and self-perception of speech and voice in men and women with idiopathic parkinson disease. Am J Speech-Lang Pathol, 6(2), 85–94. 10.1044/1058-0360.0602.85

Friston, K. (2010). The free-energy principle: a unified brain theory? Nat Rev Neurosci, 11(2), 127–138. 10.1038/nrn2787

Galvan, A., Devergnas, A., & Wichmann, T. (2015). Alterations in neuronal activity in basal ganglia-thalamocortical circuits in the parkinsonian state. Front Neuroanat, 9, 5. 10.3389/fnana.2015.00005

Garrido, M. I., Kilner, J. M., Stephan, K. E., & Friston, K. J. (2009). The mismatch negativity: a review of underlying mechanisms. Clin Neurophysiol, 120(3), 453–463. 10.1016/j.clinph.2008.11.029

Georgiev, D., Jahanshahi, M., Dreo, J., Cus, A., Pirtosek, Z., & Repovs, G. (2015). Dopaminergic medication alters auditory distractor processing in Parkinson’s disease. Acta Psychol (Amst), 156, 45–56. 10.1016/j.actpsy.2015.02.001

Gimenez-Aparisi, G., Guijarro-Estelles, E., Chornet-Lurbe, A., Ballesta-Martinez, S., Pardo-Hernandez, M., & Ye-Lin, Y. (2023). Early detection of Parkinson’s disease: Systematic analysis of the influence of the eyes on quantitative biomarkers in resting state electroencephalography. Heliyon, 9(10), e20625. 10.1016/j.heliyon.2023.e20625

Glomb, K., Cabral, J., Cattani, A., Mazzoni, A., Raj, A., & Franceschiello, B. (2022). Computational Models in Electroencephalography. Brain Topogr, 35(1), 142–161. 10.1007/s10548-021-00828-2

Groppe, D. M., Urbach, T. P., & Kutas, M. (2011). Mass univariate analysis of event-related brain potentials/fields I: a critical tutorial review. Psychophysiology, 48(12), 1711–1725. 10.1111/j.1469-8986.2011.01273.x

Han, C. X., Wang, J., Yi, G. S., & Che, Y. Q. (2013). Investigation of EEG abnormalities in the early stage of Parkinson’s disease. Cogn Neurodyn, 7(4), 351–359. 10.1007/s11571-013-9247-z

Hartoyo, A., Cadusch, P. J., Liley, D. T. J., & Hicks, D. G. (2019). Parameter estimation and identifiability in a neural population model for electro-cortical activity. PLoS Comput Biol, 15(5), e1006694. 10.1371/journal.pcbi.1006694

Hartoyo, A., Cadusch, P. J., Liley, D. T. J., & Hicks, D. G. (2020). Inferring a simple mechanism for alpha-blocking by fitting a neural population model to EEG spectra. PLoS Comput Biol, 16(4), e1007662. 10.1371/journal.pcbi.1007662

Ho, A. K., Bradshaw, J. L., Iansek, R., & Alfredson, R. (1999). Speech volume regulation in Parkinson’s disease: effects of implicit cues and explicit instructions. Neuropsychologia, 37(13), 1453–1460. 10.1016/s0028-3932(99)00067-6

Jeleazcov, C., Fechner, J., & Schwilden, H. (2005). Electroencephalogram monitoring during anesthesia with propofol and alfentanil: the impact of second order spectral analysis. Anesth Analg, 100(5), 1365–1369. 10.1213/01.ANE.0000148689.35951.BA

Jeleazcov, C., & Schwilden, H. (2003). [Bispectral analysis does not differentiate between anaesthesia EEG and a linear random process]. Biomed Tech (Berl), 48(10), 269–274. 10.1515/bmte.2003.48.10.269 (Die bispektrale Analyse unterscheidet das Narkose-EEG nicht von einem linearen Zufallsprozess.)

Kalia, L. V., & Lang, A. E. (2015). Parkinson’s disease. Lancet, 386(9996), 896–912. 10.1016/S0140-6736(14)61393-3

Lagopoulos, J., Gordon, E., Barhamali, H., Lim, C. L., Li, W. M., Clouston, P., & Morris, J. G. (1998). Dysfunctions of automatic (P300a) and controlled (P300b) processing in Parkinson’s disease. Neurol Res, 20(1), 5–10. 10.1080/01616412.1998.11740476

Liao, X. Y., Gao, Y. X., Qian, T. T., Zhou, L. H., Li, L. Q., Gong, Y., & Ye, T. F. (2024). Bibliometric analysis of electroencephalogram research in Parkinson’s disease from 2004 to 2023. Front Neurosci, 18, 1433583. 10.3389/fnins.2024.1433583

Liao, X. Y., Jiang, Y. E., Xu, R. J., Qian, T. T., Liu, S. L., & Che, Y. (2025). A bibliometric analysis of electroencephalogram research in stroke: current trends and future directions. Front Neurol, 16, 1539736. 10.3389/fneur.2025.1539736

Liley, D. T., Cadusch, P. J., & Dafilis, M. P. (2002). A spatially continuous mean field theory of electrocortical activity. Network, 13(1), 67-113. https://www.ncbi.nlm.nih.gov/pubmed/11878285

Liley, D. T. J., Cadusch, P. J., & Wright, J. J. (1999). A continuum theory of electrocortical activity. Neurocomputing, 26-27, 795–800. 10.1016/S0925-2312(98)00149-0

Liley, D. T. J., & Muthukumaraswamy, S. D. (2020). Evidence that alpha blocking is due to increases in system-level oscillatory damping not neuronal population desynchronisation. Neuroimage, 208, 116408. 10.1016/j.neuroimage.2019.116408

Liu, M., Liu, B., Ye, Z., & Wu, D. (2023). Bibliometric analysis of electroencephalogram research in mild cognitive impairment from 2005 to 2022. Front Neurosci, 17, 1128851. 10.3389/fnins.2023.1128851

McGregor, M. M., & Nelson, A. B. (2019). Circuit Mechanisms of Parkinson’s Disease. Neuron, 101(6), 1042–1056. 10.1016/j.neuron.2019.03.004

McKeown, D. J., Jones, M., Pihl, C., Finley, A. J., Kelley, N., Baumann, O., Schinazi, V. R., Moustafa, A. A., Cavanagh, J. F., & Angus, D. J. (2024). Medication-invariant resting aperiodic and periodic neural activity in Parkinson’s disease. Psychophysiology, 61(4), e14478. 10.1111/psyp.14478

Mollaei, F., Shiller, D. M., Baum, S. R., & Gracco, V. L. (2019). The Relationship Between Speech Perceptual Discrimination and Speech Production in Parkinson’s Disease. J Speech Lang Hear Res, 62(12), 4256–4268. 10.1044/2019_JSLHR-S-18-0425

Nichols, T. E., & Holmes, A. P. (2002). Nonparametric permutation tests for functional neuroimaging: a primer with examples. Hum Brain Mapp, 15(1), 1–25. 10.1002/hbm.1058

Nieto-Escamez, F., Obrero-Gaitan, E., & Cortes-Perez, I. (2023). Visual Dysfunction in Parkinson’s Disease. Brain Sci, 13(8). 10.3390/brainsci13081173

Norouzi, H., Ietswaart, M., Adair, J., & Learmonth, G. (2025). A meta-analysis of periodic and aperiodic M//EEG components in Parkinson’s disease. 10.1101/2025.07.07.663454

Oostenveld, R., Fries, P., Maris, E., & Schoffelen, J. M. (2011). FieldTrip: Open source software for advanced analysis of MEG, EEG, and invasive electrophysiological data. Comput Intell Neurosci, 2011, 156869. 10.1155/2011/156869

Ouyang, G., Hildebrandt, A., Sommer, W., & Zhou, C. (2017). Exploiting the intra-subject latency variability from single-trial event-related potentials in the P3 time range: A review and comparative evaluation of methods. Neurosci Biobehav Rev, 75, 1–21. 10.1016/j.neubiorev.2017.01.023

Ozmus, G., Yerlikaya, D., Gokceoglu, A., Emek Savas, D.D., Cakmur, R., Donmez Colakoglu, B., & Yener, G. G. (2017). Demonstration of Early Cognitive Impairment in Parkinson’s Disease with Visual P300 Responses. Noro Psikiyatr Ars, 54(1), 21–27. 10.5152/npa.2016.12455

Polverino, P., Ajcevic, M., Catalan, M., Mazzon, G., Bertolotti, C., & Manganotti, P. (2022). Brain oscillatory patterns in mild cognitive impairment due to Alzheimer’s and Parkinson’s disease: An exploratory high-density EEG study. Clin Neurophysiol, 138, 1–8. 10.1016/j.clinph.2022.01.136

Ribeiro, T. L., Jendrichovsky, P., Yu, S., Martin, D. A., Kanold, P. O., Chialvo, D. R., & Plenz, D. (2024). Trial-by-trial variability in cortical responses exhibits scaling of spatial correlations predicted from critical dynamics. Cell Rep, 43(2), 113762. 10.1016/j.celrep.2024.113762

Risacher, S. L., & Apostolova, L. G. (2023). Neuroimaging in Dementia. Continuum (Minneap Minn), 29(1), 219–254. 10.1212/con.0000000000001248

Sabbagh, M. N., Boada, M., Borson, S., Doraiswamy, P. M., Dubois, B., Ingram, J., Iwata, A., Porsteinsson, A. P., Possin, K. L., Rabinovici, G. D., Vellas, B., Chao, S., Vergallo, A., & Hampel, H. (2020). Early Detection of Mild Cognitive Impairment (MCI) in an At-Home Setting. J Prev Alzheimers Dis, 7(3), 171–178. 10.14283/jpad.2020.22

Schwilden, H., & Jeleazcov, C. (2002). Does the EEG during isoflurane/alfentanil anesthesia differ from linear random data? J Clin Monit Comput, 17(7-8), 449-457. 10.1023/a:1026284321451

Stam, C. J. (2005). Nonlinear dynamical analysis of EEG and MEG: review of an emerging field. Clin Neurophysiol, 116(10), 2266–2301. 10.1016/j.clinph.2005.06.011

Stam, C. J., Pijn, J. P., Suffczynski, P., & Lopes da Silva, F.H. (1999). Dynamics of the human alpha rhythm: evidence for non-linearity? Clin Neurophysiol, 110(10), 1801–1813. 10.1016/s1388-2457(99)00099-1

Stanzione, P., Semprini, R., Pierantozzi, M., Santilli, A. M., Fadda, L., Traversa, R., Peppe, A., & Bernardi, G. (1998). Age and stage dependency of P300 latency alterations in non-demented Parkinson’s disease patients without therapy. Electroencephalogr Clin Neurophysiol, 108(1), 80–91. 10.1016/s0168-5597(97)00070-1

Stein, R. B., Gossen, E. R., & Jones, K. E. (2005). Neuronal variability: noise or part of the signal? Nat Rev Neurosci, 6(5), 389–397. 10.1038/nrn1668

Stoffers, D., Bosboom, J. L., Deijen, J. B., Wolters, E. C., Berendse, H. W., & Stam, C. J. (2007). Slowing of oscillatory brain activity is a stable characteristic of Parkinson’s disease without dementia. Brain, 130(Pt 7), 1847-1860. 10.1093/brain/awm034

Tanaka, H., Koenig, T., Pascual-Marqui, R. D., Hirata, K., Kochi, K., & Lehmann, D. (2000). Event-related potential and EEG measures in Parkinson’s disease without and with dementia. Dement Geriatr Cogn Disord, 11(1), 39–45. 10.1159/000017212

Uslu, A., Ergen, M., Demirci, H., Lohmann, E., Hanagasi, H., & Demiralp, T. (2020). Event-related potential changes due to early-onset Parkinson’s disease in parkin (PARK2) gene mutationcarriers and non-carriers. Clin Neurophysiol, 131(7), 1444–1452. 10.1016/j.clinph.2020.02.030

Wang, L., Kuroiwa, Y., Li, M., Kamitani, T., Wang, J., Takahashi, T., Suzuki, Y., Ikegami, T., & Matsubara, S. (2000). The correlation between P300 alterations and regional cerebral blood flow in non-demented Parkinson’s disease. Neurosci Lett, 282(3), 133–136. 10.1016/s0304-3940(00)00886-7

Weil, R. S., Schrag, A. E., Warren, J. D., Crutch, S. J., Lees, A. J., & Morris, H. R. (2016). Visual dysfunction in Parkinson’s disease. Brain, 139(11), 2827–2843. 10.1093/brain/aww175

Wolff, A., Yao, L., Gomez-Pilar, J., Shoaran, M., Jiang, N., & Northoff, G. (2019). Neural variability quenching during decision-making: Neural individuality and its prestimulus complexity. Neuroimage, 192, 1–14. 10.1016/j.neuroimage.2019.02.070

Xu, H., Gu, L., Zhang, S., Wu, Y., Wei, X., Wang, C., Xu, Y., & Guo, Y. (2022). N200 and P300component changes in Parkinson’s disease: a meta-analysis. Neurol Sci, 43(12), 6719–6730. 10.1007/s10072-022-06348-6

Yassine, S., Almarouk, S., Gschwandtner, U., Auffret, M., Fuhr, P., Verin, M., & Hassan, M. (2024). Electrophysiological signatures of anxiety in Parkinson’s disease. Transl Psychiatry, 14(1), 66. 10.1038/s41398-024-02745-x

Yi, G. S., Wang, J., Deng, B., & Wei, X. L. (2017). Complexity of resting-state EEG activity in the patients with early-stage Parkinson’s disease. Cogn Neurodyn, 11(2), 147–160. 10.1007/s11571-016-9415-z

